# Repetitive induction of a hibernation-like brain state slows amyloid pathology

**DOI:** 10.64898/2025.12.26.694269

**Authors:** Ikumi Oomoto, Yong Huang, Maya Odagawa, Takeshi Sakurai, Genshiro A. Sunagawa, Hiroki Sasaguri, Masanori Murayama

## Abstract

Alzheimer’s disease (AD) is characterized by the gradual accumulation of amyloid pathology before the onset of cognitive impairment, yet effective strategies to alter disease progression remain limited. Here, we tested whether induction of a hibernation-like physiological state can slow amyloid pathology in an AD mouse model. To achieve sustained effects, we developed a protocol for repeated induction of a hypometabolic/hypothermic state and found that it significantly slowed plaque accumulation in a duration-dependent manner. This intervention also attenuated neuroinflammation and the formation of dystrophic neurites associated with plaques. Notably, this approach does not target specific molecular pathways but instead modulates global brain state. Our findings establish a proof-of-concept for a brain-state– based therapeutic paradigm for modifying disease progression in AD.

## Main Text

Alzheimer’s disease (AD) is one of the most pressing global health challenges, with the number of patients increasing rapidly due to aging populations. Despite decades of research, modifying the disease course or preventing progression remains difficult and continues to be a significant and unresolved issue. The central pathological feature in the early stages of AD is the extracellular accumulation of amyloid-β (Aβ) peptides, driven by an imbalance between Aβ production and clearance. Aβ is generated through proteolytic processing of the amyloid precursor protein (APP), and the aggregation-prone Aβ42 species accumulates to form toxic extracellular plaques (*1, 2*). While Aβ is continuously produced, age-related impairment of Aβ clearance is thought to play a major role in its accumulation, particularly in the aging brain (*3*). Notably, amyloid pathology typically precedes the onset of cognitive decline by many years, emphasizing the importance of identifying strategies to suppress or delay early Aβ deposition (*4*–*6*). Approaches capable of modifying these early pathological processes may hold substantial therapeutic potential for preventing or slowing AD progression. Accumulating evidence supports the hypothesis that Aβ production and accumulation are closely linked to neuronal activity and brain metabolic demand. Synaptic activity directly enhances extracellular Aβ release (*7, 8*), and brain regions with higher lifetime metabolic rates show preferential amyloid deposition (*9*). Therefore, reductions in neuronal activity or metabolic demand may attenuate early amyloid pathology.

Hibernation is a physiological state characterized by reduced neuronal activity and metabolic demand, enabling animals to survive harsh environmental conditions. During hibernation, animals enter a state characterized by profound decreases in body temperature and metabolic rate, along with the suppression of physiological functions (*10, 11*). These changes help minimize energy consumption during prolonged periods of resource shortages. Importantly, recent research indicates that hibernation provides potent neuroprotection, preserving neuronal structure and function despite extreme physiological stress (*12, 13*). These protective effects have stimulated interest in the concept of “hibernation-based therapies.” This concept aims to harness the key mechanisms of natural hibernation to protect the brain during disease or injury. Recent advances have revealed that activation of a specific population of hypothalamic neurons, known as QRFP-producing neurons (hereafter Q neurons), can induce a reversible, hibernation-like hypometabolic and hypothermic state, termed Q neuron-induced hypometabolic/hypothermic (QIH), in non-hibernating mammals (*14*). QIH recapitulates several features of natural hibernation, including reduced metabolic demand and lowered body temperature, without requiring a genetic or evolutionary predisposition for hibernation. Recently, the therapeutic potential of QIH in brain injury has been reported (*15*). Moreover, a hibernation-like state has been shown to slow blood epigenetic aging and extend healthspan in mice, although cortical epigenetic aging appears largely unchanged (*16*). These findings underscore the systemic effects of hibernation-like states while leaving unresolved how such states impact central nervous system pathology and brain integrity.

Given the early onset of amyloid pathology in AD and the strong neuroprotective properties of hibernation, QIH at the very earliest stage of amyloid pathology emergence might provide a unique model to examine whether hypometabolic brain states or hypothermia can modulate Aβ production, aggregation, or clearance. Here, we investigated the effect of QIH on amyloid pathology in an AD model mouse that exhibits human AD patient-like amyloid pathology at an early age (*17*). By combining repetitive chemogenetic activation of hypothalamic Q neurons with histological and quantitative analyses of Aβ deposition, we demonstrated that inducing a hibernation-like state can attenuate or delay the progression of amyloid plaque accumulation.

Our findings provide new insights into the interplay between metabolic state and AD pathology and highlight the potential of synthetic torpor as a novel therapeutic framework for modifying disease processes associated with AD.

### Generation of Q-AD Tg mice and experimental scheme of repetitive QIH

To study the effect of hibernation-like behavior on the amyloid pathology, we generate a new transgenic mouse, triple-mutant Q-AD mouse, by crossbreeding two different transgenic mice; one is the Qrfp^iCre^ mouse (*14*), in which we can induce QIH by activating Q neurons, the other is a third-generation AD model mouse (*17*), which is double-mutant of *App*^*NL-F*^ and *Psen1*^*P117L*^, and exhibits aggressive phenotype in amyloid pathology (Fig. 1A). We note that the triple-mutant mice used in this study were heterozygous for Qrfp^iCre^ and homozygous for other two genes. Q-AD mice exhibit amyloid pathology at a very early age, starting from the cortex at 6 weeks old (Fig. 1B–D, Supplementary Fig. 1), similar to the original *App*^*NL-F*^*Psen1*^*P117L*^ mice (*17*). In the early stage (6-8 weeks), small Aβ plaques begin to appear in the deep layers of the lateral cortex. As the mice aged, plaque size gradually increased and became visible throughout the cortex. Since amyloid pathology was clearly observed in the somatosensory cortex from the earliest stages, we primarily quantified Aβ deposition in this region.

**Fig. 1.**
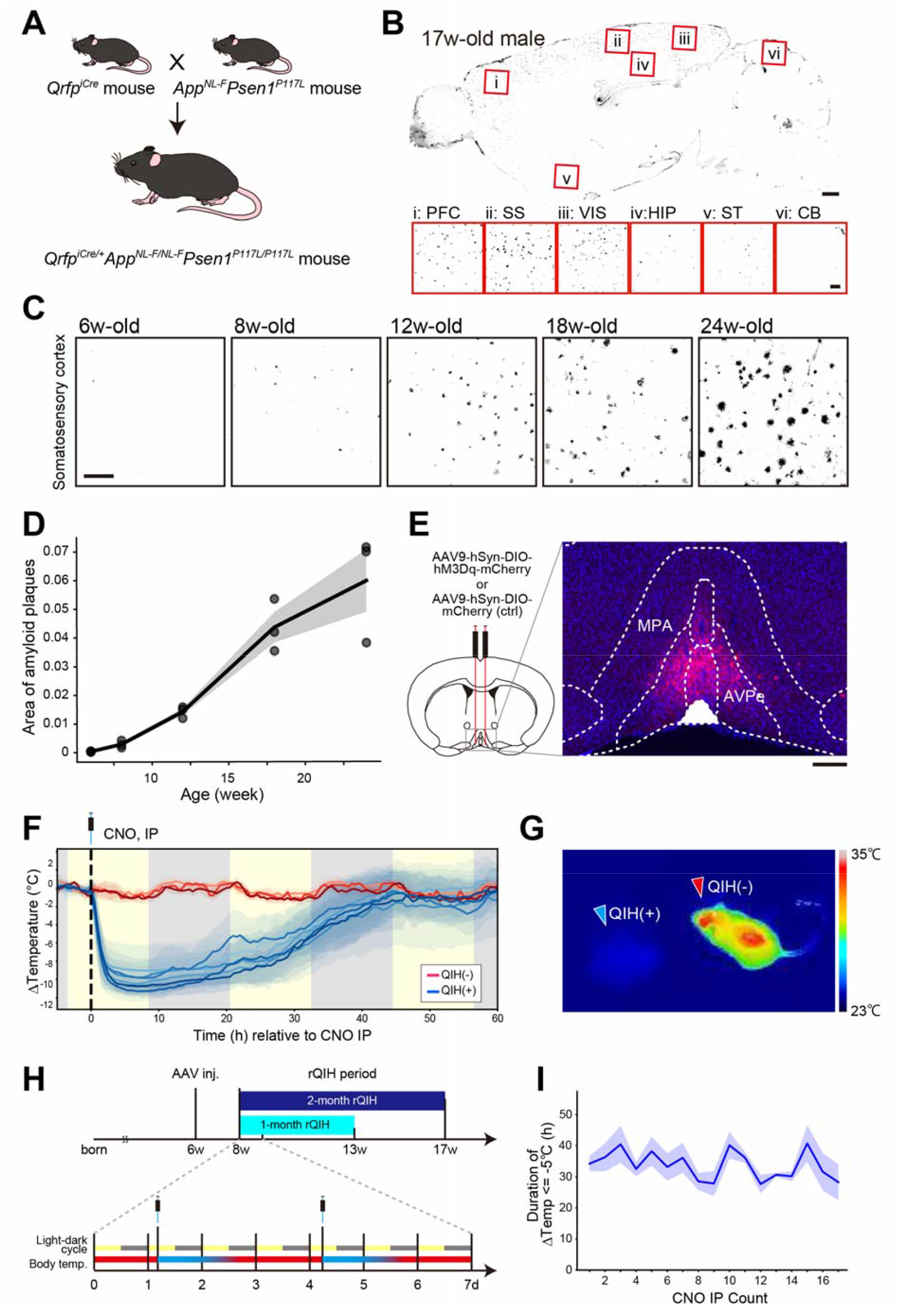
Generation of Q-AD Transgenic mice and experimental timeline of rQIH. (A) Schematic illustration of the generation of Q-AD mice. (B) Sagittal brain section showing Aβ deposition visualized by immunostaining with the 82E1 antibody in 17-week-old male Q-AD mice. Insets: higher-magnification views of the indicated region. Scale bars, 500 μm and 100 μm (inset). (C) Age-dependent progression of amyloid plaque accumulation in the somatosensory cortex of male Q-AD mice. Scale bar, 100 μm. (D) Quantification of Aβ deposition in the motor and somatosensory cortex stained with the 82E1 antibody. Thick line indicates the mean, and shaded area represents the SEM (n = 3 mice per age group). (E) Representative brain section of an AAV-injected Q-AD mouse. AAV9-hSyn-DIO-hM3Dq-mCherry (QIH(+)) or AAV9-hSyn-DIO-mCherry (QIH(-)) was injected into the AVPe/MPA. Scale bar, 100 μm. (F) Temporal profile of body temperature following DREADD-mediated activation of Q neurons in AAV-injected Q-AD mice. Body temperature traces from multiple CNO administrations in each animal are aligned to the time of intraperitoneal injection (time 0). Thick lines indicate the mean, and shaded areas represent the SD. (G) Infrared thermal image showing surface body temperature of CNO-treated Q-AD mice, acquired 3 hours after CNO administration. (H) Experimental schema of rQIH. Intraperitoneal (IP) injection of CNO was performed to chemogenetically activate hM3Dq-expressing Q neurons twice per week for 5 or 9 weeks. (I) Quantification of the duration of a ≥5°C decrease in body temperature induced by rQIH (n = 7 mice). Thick lines indicate the mean, and shaded areas represent the SEM.

To chemogenetically activate Q neurons, which leads to QIH in Q-AD mice by intraperitoneal (IP) injection of clozapine-N-oxide (CNO) (a DREADD agonist), we expressed hM3Dq– mCherry in AVPe (Anteroventral periventricular nucleus) and MPA (Medial preoptic area) by injecting 6-week-old Q-AD mice with AAV9-hSyn-DIO-hM3Dq-mCherry (Fig. 1E). The IP administration of CNO immediately induced hypothermia in QIH(+) mice, lowering their body temperature to room temperature levels (Fig. 1F, G). On average, QIH sustained around 24-36 hours upon single CNO administration in our protocol (Fig. 1F). Because mice are not natural hibernators, they do not undergo pre-hibernation metabolic adaptation or energy storage; consequently, prolonged QIH induction may lead to negative energy balance and body weight loss, imposing an excessive metabolic burden and requiring recovery periods to allow physiological re-equilibration following sustained hypometabolic states. To achieve prolonged QIH within a less-harmful range for mice, we employed a repetitive QIH (rQIH) paradigm, in which QIH was induced twice per week for either 1 month or 2 months, so that QIH could be induced for about half of the rQIH period (Fig. 1H). Repeating administration of CNO did not affect the duration of the QIH (Fig. 1I), but induced a transient decrease in body weight during the early phase of rQIH (Supplementary Fig. S2).

### rQIH Suppresses Amyloid Plaque Accumulation

We next assessed the effects of rQIH on amyloid pathology in the Q-AD mice. Immunohistochemical analyses were performed as previously described (*17, 18*), to evaluate amyloid pathology in the brains of 3-month-old Q-AD mice with a 1-month rQIH period (Fig. 2A–C) and 4-month-old Q-AD mice with a 2-month rQIH (Fig. 2D–F). The 82E1 antibody, which recognizes the N-terminal epitope of β-secretase–derived Aβ (*19*), was used to visualize dense amyloid deposits. Compared with QIH(-) control mice, in which QIH is not induced due to the absence of DREADD receptor expression, amyloid pathology was significantly attenuated by rQIH in the cortex in QIH(+) mice (Fig. 2G, I, K). The reduction in plaque burden was more pronounced in the medial cortical regions than in the lateral regions (Fig. 2B, E). Quantitative analysis of the plaques (see Methods) revealed a significant decrease in the number and total area of large (>100 μm^2^) 82E1-positive amyloid plaques, indicating an overall reduction in Aβ deposition (Fig. 2G, I; Supplementary Fig. 3). Layer-resolved analyses were performed across the major cortical layers (L1, L2/3, L4, L5, and L6; see Methods and Supplementary Fig. 1). In the 1-month rQIH paradigm, linear mixed-effects modeling detected modest layer-dependent effects, with relatively larger reductions in Aβ deposition in middle cortical layers, particularly L4 and L5, compared with superficial and deep layers (Supplementary Fig. 4). In contrast, these layer-dependent interactions were no longer evident in the 2-month rQIH group (Supplementary Fig. 4), indicating that prolonged QIH induces a more uniform suppression of amyloid pathology across cortical layers. A significant negative association was observed between plaque area and the average duration of rQIH (Fig. 2H, J; Spearman’s rank correlation, ρ = −0.744, p = 0.0023 for 1-month rQIH, and ρ = −0.799, p = 0.0018 for 2-month rQIH), indicating that rQIH is associated with a reduction in amyloid pathology in QIH(+) mice. Consistent effects were also observed in female mice (Supplementary Fig. 5).

**Fig. 2.**
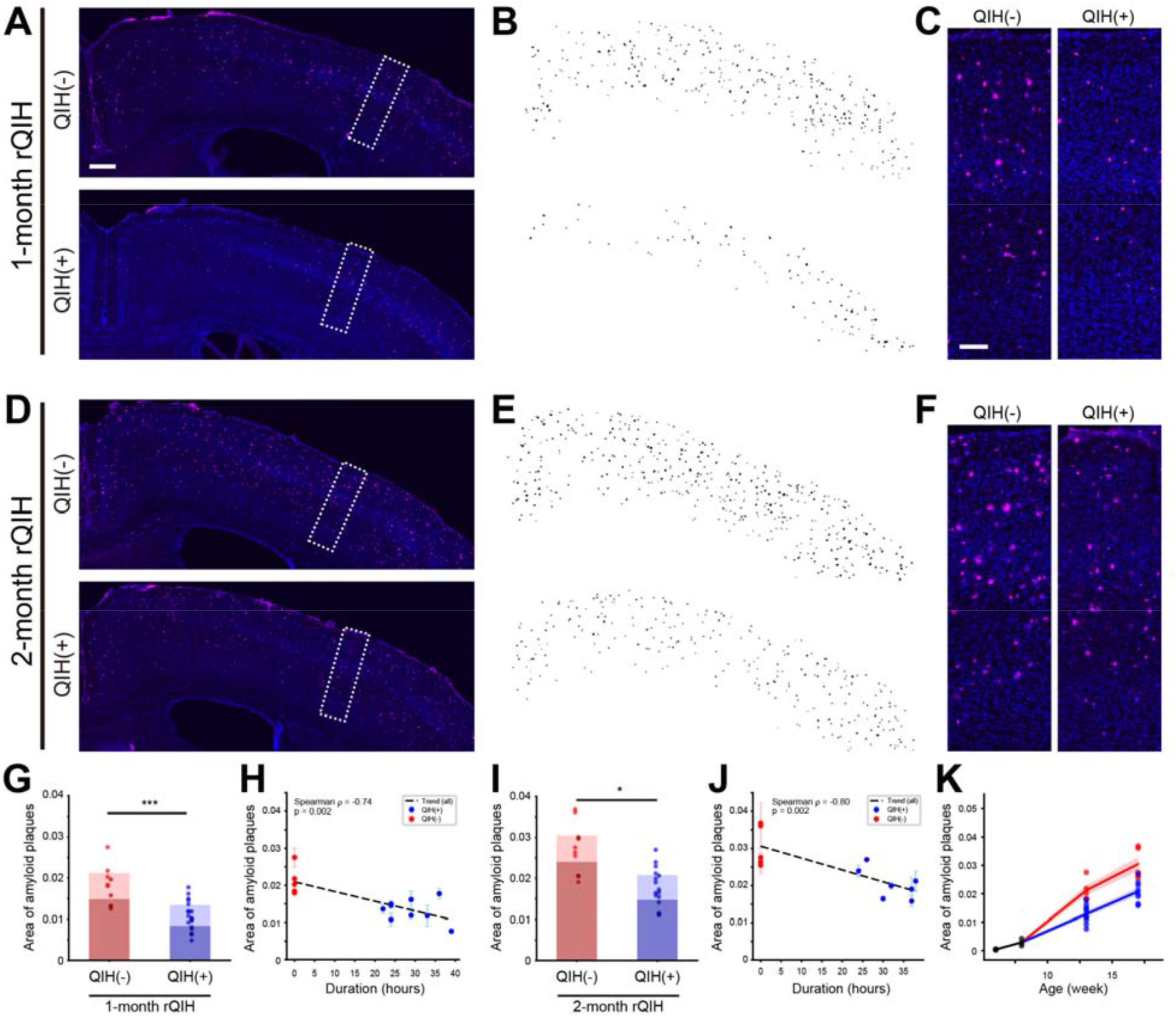
rQIH suppresses the progression of amyloid pathology. (A, D) Representative coronal brain images showing Aβ deposition visualized by immunostaining with 82E1 antibody in 1-month (A) or 2-month (D) rQIH conducted mice. Scale bar, 200 μm. (B, E) Binalized images of Aβ deposition for 1-month (B) or 2-month (E) rQIH. Plaques larger than 100 μm^2^ are visualized. (C, F) Enlarged images of the white-dashed squares in panel A or D. Scale bar, 50 μm. (G, I) Quantification of Aβ deposition in male Q-AD mice subjected to 1-month rQIH (G; QIH(+), n = 9; QIH(-), n = 5) or 2-month rQIH (I; QIH(+), n = 7; QIH(-), n = 5). Dark-colored bars indicate quantification of large plaques (>100 μm^2^). For panel G, p = 0.000999 for plaques ≥10 μm^2^ and p = 0.000999 for plaques ≥100 μm^2^. For panel H, p = 0.0101 for plaques ≥10 μm^2^ and p = 0.01768 for plaques ≥100 μm^2^ (Mann–Whitney U test). (H, J) Scatter plot showing the relationship between QIH duration and mean plaque area in the cortex of 1-month rQIH (H) or 2-month rQIH (J). Each marker represents an individual mouse; vertical bars indicate mean ± SD. QIH(-) mice were plotted at duration = 0. (K) Summary plot of amyloid plaque burden across age. The area of amyloid plaques in the cortex is plotted against age (weeks) for QIH(+) and QIH(-) mice. Data from both 1-month and 2-month rQIH cohorts are combined. Each marker represents an individual mouse; thick lines indicate the mean, and the shaded area indicates the SEM.

**Fig. 2.**
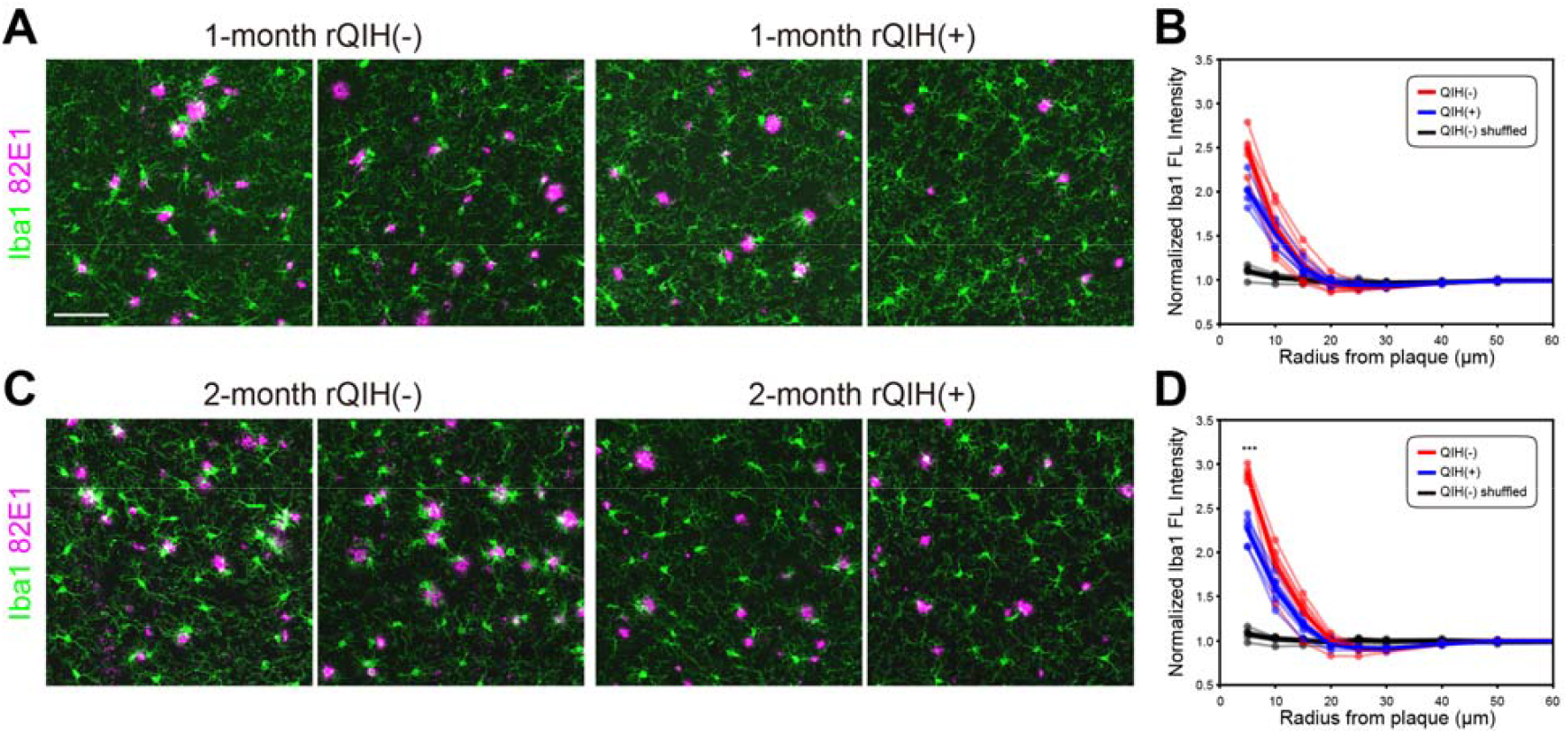
Neuroinflammation in Q-AD mice upon rQIH. (A, C) Representative coronal brain sections of mice subjected to 1-month rQIH (A) or 2-month rQIH (C), showing Aβ deposition and microglia visualized by immunostaining with 82E1 (magenta) and a microglial marker Iba1 (green) antibodies. Arrowheads indicate the amoeboid-shaped Iba1□ microglia. Scale bar, 100 μm (shown in A). (B, D) Quantification of Iba1 fluorescence intensity as a function of distance from amyloid plaques. Concentric circular regions of interest (ROIs) were drawn centered on individual plaques, and Iba1 signal intensity was measured radially from the plaque center. The x-axis represents the distance from the plaque center (μm). Red, QIH(-); blue, QIH(+). As a spatial control, shuffled ROIs generated in plaque-free regions from QIH(-) images are shown in black. Thin lines represent individual animals, thick lines represent the mean, and shaded areas indicate SEM. (B: n = 5 for QIH(−), n = 5 for QIH(+); D: n = 5 for QIH(−), n = 6 for QIH(+)). Statistical comparisons were performed between QIH(−) and QIH(+) groups only using Welch’s t-test with Benjamini–Hochberg FDR correction. ***: p_FDR = 2.18 × 10□□ at 0–5 μm; no other distance bins were significant.

**Fig. 2.**
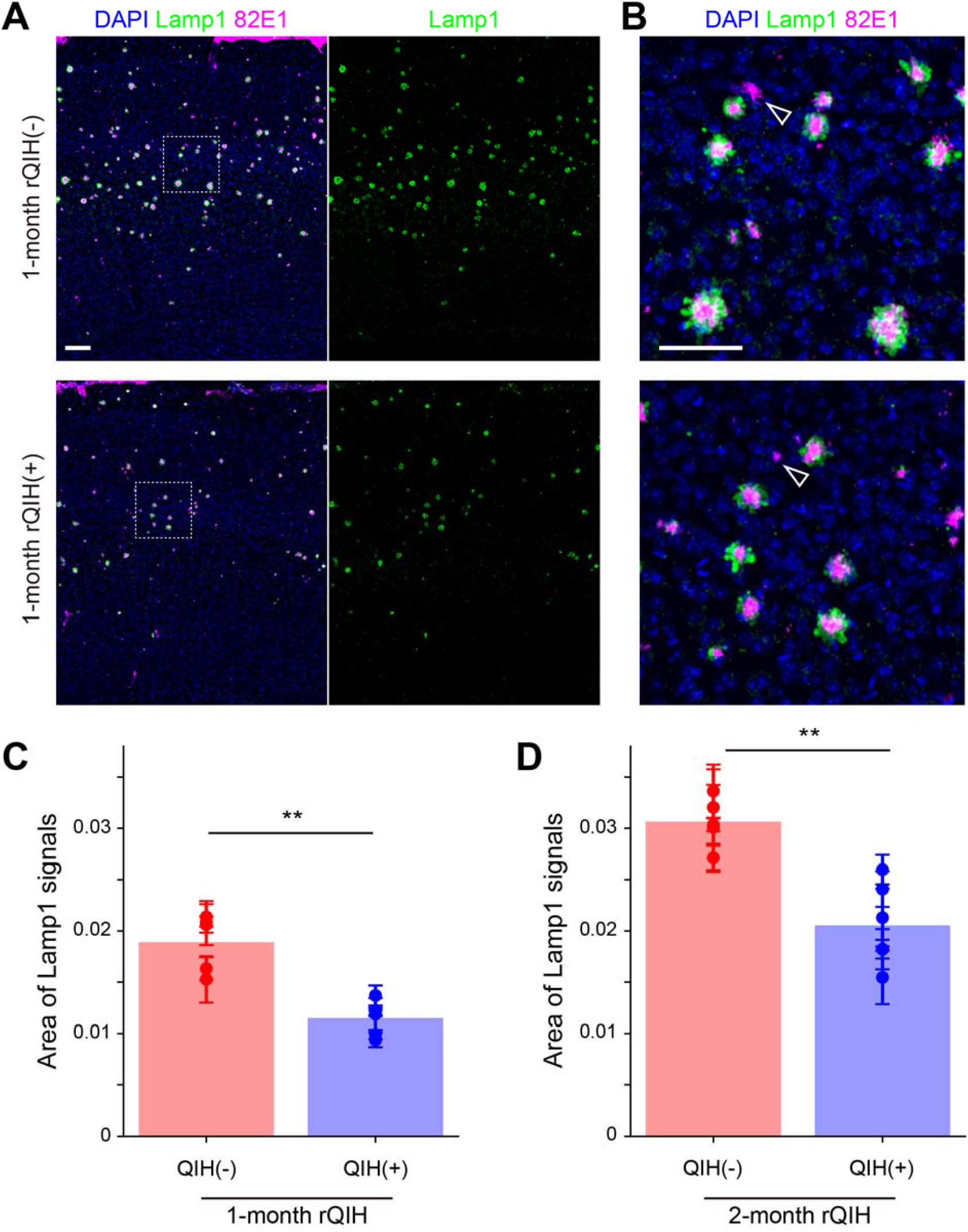
Neuritic alterations in Q-AD mice upon rQIH. (A) Coronal brain image showing Aβ deposition and neuritic alterations visualized by immunostaining with 82E1 (magenta) and dystrophic neurite marker Lamp1 (green) antibodies. Scale bar, 100 μm. (B) Enlarged image of the dashed square area in A. Arrowheads indicate the 82E1 signals lacking Lamp1 signals. Scale bar, 50 μm. (C, D) Quantification of Lamp1 signals in the cortex. For 1-month rQIH, n = 6 for QIH(+), n = 5 for QIH(-) (C; p = 0.00433, Mann–Whitney U test). For 2-month rQIH, n = 6 for QIH(+), n = 5 for QIH(-) (D; p = 0.00433, Mann–Whitney U test).

To further assess the impact of rQIH on amyloid pathology, we compared amyloid burden between mice subjected to 1 month of rQIH(−) and those subjected to 2 months of rQIH(+) (Supplementary Fig. 6). No significant differences were observed in overall Aβ deposition, layer-resolved distribution, or plaque size distribution between these conditions. These results suggest that, under the conditions tested, the present rQIH protocol can delay the progression of Aβ pathology by about one month in the present Q-AD model.

### Effects of rQIH on neuroinflammations and neuritic alterations

As amyloid pathology progresses, additional cellular and neuritic alterations accompany plaque formation (*20, 21*). We therefore examined whether rQIH-induced changes in amyloid burden were associated with modifications in these downstream features. Neuroinflammation is driven by the activation of microglia and astrocytes and is frequently observed in the brains of patients with AD and in AD model mice, along with preceding amyloid pathology (*22, 23*). To quantify plaque-associated microglial responses, concentric circular regions of interest (ROIs) were drawn centered on individual 82E1-positive plaques, and immunoreactivity for Iba1 (a microglial marker) was measured as a function of distance from the plaque center. In both QIH(+) and QIH(-) groups, Iba1 signal intensity was elevated in plaque-proximal regions relative to distal regions, indicating microglial accumulation around plaques (Fig. 3). After 1 month of rQIH, plaque-proximal Iba1 intensity in QIH(+) mice showed a decreasing trend compared to QIH(-) mice, although this difference did not reach statistical significance (Fig. 3A, B). In contrast, after 2 months of rQIH, plaque-proximal Iba1 intensity was significantly reduced in the QIH(+) group (Fig. 3C, D). These results suggest that the effect of rQIH on microglial accumulation becomes more evident at later stages of amyloid pathology. As a spatial control, concentric ROIs were generated in plaque-free regions, where Iba1 intensity remained relatively flat across distances and showed no differences between groups at any radius (Fig. 3B, D; see Methods and Supplementary Fig. 7). In addition, Iba1 signals in plaque-centered ROIs were consistently higher than in shuffled controls, confirming that microglial accumulation was specifically associated with plaques.

In addition to neuroinflammatory responses, neuritic alterations represent another hallmark that emerges during amyloid plaque progression (*24*). To examine whether rQIH influences plaque-associated neurite pathology, we evaluated dystrophic neurite (DN) formation by co-immunostaining with antibodies against 82E1 and Lamp1, a marker for lysosome-endosome vesicles (Fig. 4A, B). Lamp1-positive signals were frequently observed surrounding 82E1-positive amyloid plaques, consistent with plaque-associated neuritic dystrophy. In contrast, smaller Aβ plaques were often not accompanied by detectable Lamp1 signals (Fig. 4B). Quantitative analysis revealed that the total area of Lamp1-labeled DNs was significantly reduced in the rQIH(+) group compared with rQIH(-) controls at both 1 month and 2 months (Fig. 4C, D; p = 0.00433 for both). Together, these results indicate that rQIH reduces plaque-proximal Iba1 intensity in a stage-dependent manner, while consistently attenuating neuritic dystrophy, suggesting a coordinated attenuation of plaque-associated neuroinflammation and secondary neuritic alterations accompanying amyloid plaque progression.

## Discussion

### Summary

In this study, we generated a novel hibernation-like state–inducible AD model mouse by cross-mating an AD model line with hibernation-like state–inducible QrfpiCre mice, thereby enabling induction of QIH in an AD background (Fig. 1). Using these animals, we demonstrated that rQIH suppresses the progression of amyloid pathology (Fig. 2). The suppressive effect was greater when QIH lasted longer, and rQIH specifically reduced large 82E1-positive amyloid plaques (Fig. 2), accompanied by a reduction in neuroinflammation and dystrophic neurite formation (Fig. 3 and 4). We further established a protocol for repetitive induction of QIH by repeated administration of CNO, twice per week for one to two months. Under the experimental conditions tested, this protocol induced a hibernation-like state for prolonged periods without detectable adverse effects on animal health, as assessed by body weight, survival, and general behavioral observations.

### Proof of concept: suppressing amyloid pathology by manipulating a brain-wide physiological state

Our findings provide a proof of concept that repeated induction of a hibernation-like state can suppress the progression of early amyloid pathology by manipulating a brain-wide physiological state. Whereas amyloid pathology has most often been discussed within a framework of targeting individual molecular components of amyloid metabolism (*1, 25, 26*), our study was designed not to test a single predefined molecular pathway, but rather to ask whether a hibernation-like, integrated hypometabolic state itself can influence amyloid pathology. Although we do not yet delineate the precise molecular mechanisms underlying this effect, our results demonstrate the feasibility of modulating amyloid pathology by manipulating the global brain metabolic state.

### Mechanisms underlying the suppressive effect of QIH on amyloid pathology

Several non-mutually exclusive mechanisms could account for the observed suppression of amyloid progression during QIH. First, QIH-induced hypometabolism may reduce Aβ production and/or release. Hypometabolic states are known to broadly suppress transcriptional and translational processes (*27, 28*). Thus, QIH may downregulate APP expression and thereby reduce the supply of its cleavage product, amyloid-β (Aβ), attenuating initiation and progression of amyloid pathology. Consistent with this idea, modest down-regulation of APP expression in transgenic mouse models decreases soluble and insoluble Aβ levels and reduces amyloid plaque burden (*29*). In addition, metabolic depression may reduce neuronal activity, and if extracellular Aβ release is activity-dependent, reduced activity could decrease synaptic Aβ release and slow amyloid progression (*7, 8*). This interpretation is also consistent with reports that global metabolic downregulation, such as caloric restriction, attenuates Aβ generation and amyloid plaque deposition in AD model mice (*30*). This interpretation is also consistent with our observation that amyloid accumulation appears to be delayed during periods of rQIH (Supplementary Fig. 6), raising the possibility that intermittent hypometabolic states cumulatively slow amyloid progression.

Second, QIH may enhance extracellular Aβ clearance. The glymphatic system is highly active during sleep and facilitates the removal of extracellular metabolites, including Aβ (*31, 32*). Moreover, accumulating evidence indicates that sleep and circadian changes are not merely secondary to neurodegeneration but can actively regulate neurodegenerative processes, including Aβ metabolism (*33*). If the QIH-induced state engages sleep-like pathways—through reduced metabolism, altered perfusion, or changes in fluid dynamics—the reduction in Aβ burden observed here could reflect, at least in part, enhanced clearance rather than solely decreased production.

Third, QIH may dampen neuroinflammation, potentially mitigating immune-mediated amplification of AD pathology. In Alzheimer’s disease, chronic microglial activation, complement dysregulation, and broader immune-driven neuroinflammation are widely implicated in amplifying pathology and neurodegeneration (*22, 34*–*36*). In natural hibernators, peripheral immune cell numbers and immune responses are reduced in a reversible manner (*37, 38*). In addition, hibernation is associated with changes in microglial morphology and distribution in the CNS and may shift the brain toward a less inflammatory state (*13, 39*). Thus, rQIH might attenuate immune-dependent neuronal injury around Aβ deposits, contributing to the observed reduction in dystrophic neurite formation.

Fourth, environmental cold exposure during the QIH protocol may contribute to the observed effects. However, the contribution of temperature per se is difficult to interpret, because ambient cooling can increase metabolic demand as mammals attempt to maintain body temperature (*40*). Notably, mild or chronic exposure to environmental cold has been reported not to significantly alter Aβ or tau pathology (*41*), suggesting that reduced ambient temperature per se may be insufficient to robustly modulate amyloid-related processes. Therefore, the suppressive effect observed here is unlikely to be explained by environmental cold exposure per se (i.e., a decrease in ambient temperature) alone. Rather, it is more likely that one or more of the mechanisms proposed above (i.e., reduced Aβ production, enhanced clearance, and dampened neuroinflammation) may act in combination, potentially synergistically, within the coordinated hypometabolic/hypothermic state induced by QIH.

### Physiological and therapeutic implications

The suppressive effect on amyloid progression observed here can be interpreted as a slowing, or temporary arrest, of pathological advancement. Such an effect could preserve a therapeutic time window, thereby enabling subsequent and potentially more interventional treatment strategies. Importantly, our approach leverages endogenous physiological mechanisms rather than directly targeting pathology through a single pharmacological pathway. If QIH influences amyloid pathology in part by enhancing clearance of extracellular metabolites, similar clearance-associated benefits may extend to other age-related neurodegenerative diseases (*42, 43*). Notably, the effects of rQIH on microglial responses appeared to be stage-dependent, with clearer modulation observed at later stages of pathology (Fig. 3). This may reflect the relatively limited plaque-associated enrichment of microglia at early stages of AD pathology (e.g., at 13 weeks of age), resulting in a reduced spatial contrast in Iba1 signal intensity relative to plaque proximity, rather than indicating a lack of responsiveness to rQIH. This interpretation is consistent with previous studies showing that microglial recruitment to amyloid plaques is progressive and becomes more pronounced as pathology advances (*44*). Taken together, inducing a controlled hypometabolic brain state, or mimicking key molecular features of hibernation, may represent a novel and promising avenue for disease-modifying strategies in preclinical or early-stage AD.

### Limitations and future directions

This proof-of-concept study has several limitations. First, our analyses focused on early-stage amyloid pathology; the long-term effects following rQIH, the impact of later-stage rQIH, and effects on cognitive function remain unexplored, as behavioral assessments were not performed in the present study. Second, while we primarily examined Aβ pathology, other major disease axes, including tau pathology, were not investigated. Third, because Aβ production is significantly increased in the AD model mice we used due to the combined effects of multiple mutations, we cannot determine the extent to which rQIH affects Aβ production at more intrinsic levels. In addition, the double-mutant model used here exhibits extremely rapid progression and a severe phenotype; studies in milder models may reveal larger effects, potentially extending beyond suppressing progression to producing measurable improvement.

A key open question is which processes—reduced Aβ production/release, enhanced extracellular clearance, and/or immune modulation—are necessary and sufficient for the suppressive effect observed during QIH. Disentangling these contributions will require direct measurements of soluble and insoluble Aβ species, analyses of APP processing pathways (e.g., western blot, proteome analysis), and tests of clearance mechanisms (including glymphatic function), along with profiling of neuroinflammatory and immune-related responses.

Another important issue is whether the suppressive effect arises primarily from hypothermia or metabolic suppression. Prior work shows that manipulating thermoregulatory circuits can induce metabolic suppression and hibernation-like states even under warm ambient conditions (*45*). Such approaches could help test the effect of metabolic suppression while minimizing hypothermia, thereby clarifying their relative contributions. Conversely, because our mice were maintained at room temperature, the reduction in body temperature was limited to room-temperature levels; deeper hypothermia—more typical of natural hibernation—might be achieved by maintaining animals in a cold room during QIH, potentially enhancing suppression.

## Supporting information

Supplementary data (including materials and methods, supplementary figures)

## Acknowledgments

We thank J. Nagai and all laboratory members for valuable comments and discussions on the project. We also thank the Support Unit for Bio-Material Analysis, RIKEN CBS Research Resources Division, particularly R. Ando for technical assistance with AAV injections, A. Ito for technical assistance with embryo manipulation, and Y. Mizukami, N. Kaieda, and K. Ueno for animal care. During the preparation of this work, the authors used ChatGPT and Grammarly to improve language and readability. The authors reviewed and edited the content as needed and take full responsibility for the content of the publication.

## Funding

Japan Society for the Promotion of Science KAKENHI JP18K07402 (HS)

Japan Society for the Promotion of Science KAKENHI JP23H04941 (GAS)

Japan Society for the Promotion of Science KAKENHI JP24H02313 (MM)

Japan Science and Technology Agency FOREST Program JPMJFR2066 (GAS)

Japan Agency for Medical Research and Development Brain/MINDS 2.0 Project JP23wm0625001 (HS, MM)

## Author contributions

Conceptualization: IO, HS, GAS, MM

Methodology: IO, HS, GAS, MM

Investigation: IO, HY, MO

Visualization: IO, MM

Funding acquisition: HS, GAS, MM

Project administration: IO, MM

Supervision: HS, GAS, MM

Writing – original draft: IO, MM

Writing – review & editing: HY, HS, GAS, TS, MM

## Competing interests

The authors declare that they have no competing interests.

## Data, code, and materials availability

The data supporting the findings of this study and the custom code used for data analysis are available from the corresponding author upon reasonable request. Representative datasets and analysis scripts will be made publicly available in a permanent repository upon publication. Materials generated in this study are available from the corresponding author upon reasonable request.

## Supplementary Materials

Materials and Methods

Figs. S1 to S7

References (*46*)

## Notes

### Competing Interest Statement

The authors have declared no competing interest.

### Summary of Updates

Figure 3 added (original Figure 3 renumbered as Figure 4). Figures and supplementary figures updated. Text revised for clarity.

