## Supplementary data (including materials and methods, supplementary figures) for "Repetitive induction of a hibernation-like brain state slows amyloid pathology"

Science
AAAS

### Supplementary Materials for

#### Repetitive induction of a hibernation-like brain state slows amyloid pathology

**Authors:** Ikumi Oomoto[1], Yong Huang[1], Maya Odagawa[1], Takeshi Sakurai[2,3,4], Genshiro A. Sunagawa[5*], Hiroki Sasaguri[6*], Masanori Murayama[1*]

**The PDF file includes:**

Materials and Methods
Figs. S1 to S7
References

#### Materials and Methods

Animals

All animal experiments were performed in accordance with the institutional guidelines and were approved by the Animal Experiment Committee at RIKEN. Double-mutant AD model mice (App/Psen1; (*17*)) and $Qrfp^{iCre}$ mice (*14*) were used. The triple transgenic Q-AD mouse was generated by crossing App/Psen1 mice with Qrfp mice or through embryonic manipulation. Both male and female mice aged 6-24 weeks and weighing over 16.0 g for males and 15.0 g for females at 6 weeks old were used in the study without disparity. Mice were housed under controlled conditions (22–24 °C, 40–60% humidity, 12 h light/12 h dark light cycle) with ad libitum access to food and water.

Viruses

Adeno-associated viruses (AAV) were purchased from Addgene: AAV9-hSyn-DIO-hM3D(Gq)-mCherry (44361-AAV9) and AAV9-hSyn-DIO-mCherry (50459-AAV9).

Surgery

For AAV vector injection, 6-week-old mice were anesthetized with isoflurane (2%) for induction. Once the mice failed to respond to stimuli, we administered a hypodermic injection of the combination agent (*46*) – Medetomidine/Midazolam/Butorphanol (MMB) – at a dose of 5 mL/kg body weight for the procedure. We dispensed MMB solution with medetomidine (0.12 mg/kg body weight), midazolam (0.32 mg/kg body weight), butorphanol (0.4 mg/kg body weight), and saline. After anesthesia with MMB, the head hair of the mice was shaved, and they were placed in head holders (SG-4N, NARISHIGE). During surgery, the mice's body temperature was maintained at 36–37 °C using a feedback-controlled heat pad (BWT-100, Bio Research Center), and their eyes were coated with ointment (Neo-Medrol EE Ointment, Pfizer Inc.). The scalp was cleaned with 70% ethanol and iodine, and then incised to expose the skull surface. Mice underwent injection of AAV9-hSyn-DIO-hM3Dq-mCherry or AAV9-hSyn-DIO-mCherry into the AVPe/MPA: anterior–posterior (AP), +0.83 mm; medial–lateral (ML), ± 0.3 mm; dorsal–ventral (DV), −5.25 mm; 0.1 μl in each site at a controlled rate of 0.1 μl per 10 min using a pulled fine-glass capillary. The needle was kept in place for 10 min before and after the injection. After the surgery, the mice were dispensed a medetomidine-reversing agent, atipamezole hydrochloride (ANTISEDAN, zoetis Inc.) solution at a dose of 0.12 mg/kg body weight, then the mice were recovered on a heating pad. The waiting period for recovery and virus expression for the experiments was 2 weeks.

For the implantation of a small motion measurement device (nano tag; KISSEI COMTEC), mice were anesthetized with isoflurane (2%) for induction, followed by MMB. The hair on the back was shaved, and the skin was cleaned with 70% ethanol and iodine. After making a vertical incision approximately 1.5 cm in length in the skin, the nanotag was inserted, and the skin was sutured with nylon thread (ER2004NA45-KF2, alfresa). The suture site was bonded using the medical adhesive derma+flex (CHEMENCE) to maintain joint strength. After the surgery, the mice were dispensed atipamezole hydrochloride.

Chemogenetic induction of QIH

CNO (BML-NS105-0025, Enzo) was dissolved in saline at a dose of 1mg ml$^{-1}$ and frozen at −20 °C. The CNO solution was thawed, diluted 10-fold with saline (100 µg ml$^{-1}$), and administered intraperitoneally at a dose of 1 mg kg−1 to mice.

Body temperature and locomotion recordings

Body temperature and locomotor activity were monitored at 5-minute intervals using an implanted nano tag. The acquired data were imported into the accompanying software (nanotag Viewer, KISSEI COMTEC) and analyzed using custom Python scripts.

For thermographic analysis, mice were placed in experimental cages (28.5 × 17.5 × 12.5 cm) and imaged with an infrared thermal camera (Mini2 V2, HIKMICRO) mounted on an iPhone 13 mini (Apple), positioned 30 cm above the cage floor.

Immunohistochemistry, imaging, and analysis

The mice were deeply anesthetized by intraperitoneal injection of MMB and perfused transcardially with PBS supplemented with heparin (10 units/mL), followed by 4% paraformaldehyde (PFA) in phosphate-buffer saline (PBS; pH 7.4). The brains were then harvested and post-fixated in PFA overnight at 4 °C. The brains were cryoprotected with 30% sucrose in PBS and sectioned into 40-µm-thick coronal or sagittal sections using a freezing microtome (ROM-380, Yamato).

For immunohistochemistry, the sections were incubated in a blocking solution (2% normal goat serum and 0.3% TritonX-100 in PBS) for 30 min at room temperature. The sections were then incubated in a blocking solution containing a primary antibody, overnight at 4 °C, and subsequently washed three times with a washing buffer (0.3% TritonX-100 in PBS). The sections were incubated in a blocking solution containing a secondary antibody conjugated to a fluorescent dye for 2 h at room temperature, then washed three times with the washing buffer. The sections were stained for 15 min with DAPI (D9542, sigma) diluted in PBS and then mounted onto slides using a Fluoromount/Plus anti-fading agent (K048; Diagnostic BioSystems). Section images were obtained using a confocal laser-scanning microscope (FV-3000, Evident). Quantification of immunofluorescent signals was performed using ImageJ software.

Primary and secondary antibodies used in this study are as follows: anti-Iba1 (WAKO, 019-19741), anti-human amyloid β (N; 82E1; IBL, #10323), anti-LAMP1 (Abcam, ab208943), Goat anti-Mouse IgG (H+L) Highly Cross-Adsorbed Secondary Antibody, Alexa Fluor Plus 647 (Thermo, A32728), and Goat anti-Rabbit IgG (H+L) Highly Cross-Adsorbed Secondary Antibody, Alexa Fluor Plus 488 (Thermo, A32731).

Quantification and statistical analysis

Image quantification was performed using ImageJ/Fiji (NIH). For amyloid plaque size quantification, immunofluorescence images of 82E1-stained coronal brain sections were binarized using the MaxEntropy thresholding method, applied consistently across samples within each experiment. To minimize the inclusion of noise and non-specific signals, particles smaller than 10 µm$^2$ were excluded from all analyses. Plaque-related analyses were conducted on particles ≥10 µm$^2$, and analyses focusing on large plaques applied an additional threshold of ≥100 µm$^2$. For dystrophic neurite quantification, Lamp1 immunofluorescence images were subjected to background subtraction (rolling ball radius: 50 pixels), followed by binarization using the Moments thresholding method, applied consistently across samples within each experiment. To further reduce noise and non-specific signals, particles smaller than 50 µm$^2$ were excluded from all analyses. For quantification of inflammation, Iba1 immunofluorescence images were processed using background subtraction (rolling ball radius: 50 pixels) and Otsu thresholding.

Quantification was performed in two ways: (i) across the entire cortical region and (ii) separately within individual cortical layers (L1, L2/3, L4, L5, and L6). Cortical layers were manually defined as regions of interest (ROIs) based on DAPI nuclear staining as anatomical landmarks.

In addition, plaque-centered radial analyses were performed using a custom ImageJ macro to quantify microglial signals as a function of distance from individual plaques. Plaque centroids were identified after MaxEntropy-based segmentation, and concentric ring ROIs (5, 10, 15, 20, 25, 30, 40, 50, and 100 μm) were generated. Plaques located within 100 μm of image borders were excluded to ensure complete ring sampling. For each ring, raw integrated intensity was measured, and normalized intensity per ring area were calculated. Plaque-level values were averaged per mouse to obtain one value per ring for statistical analysis. To control for potential inter-session baseline differences, radial intensity values were normalized to the distal 50–100 μm ring, which showed no group difference and was used as a reference.

For whole-cortex quantification, statistical comparisons between control (QIH(-)) and QIH(+) groups were performed using the two-sided Mann–Whitney U test. Correlations between QIH duration and plaque size were assessed using Spearman's rank correlation coefficient.

Layer-resolved analyses were performed using a linear mixed-effects model (LMM). Condition (QIH(-) vs QIH(+)), Layer (L1–L6), and their interaction were treated as fixed effects, while mouse identity was included as a random effect to account for repeated measurements across cortical layers within the same animal.

For radial analyses, group differences at each ring distance were assessed using mouse-level comparisons with two-sided Mann-Whitney U tests, followed by multiple-comparison correction using the Benjamini–Hochberg procedure to control the false discovery rate (FDR).

All statistical analyses were performed using Python (v3.7.11), primarily utilizing the statsmodels and SciPy libraries. Data are presented as mean ± SEM unless otherwise specified. Exact p-values are reported in the figures or figure legends. Statistical significance was defined as $p < 0.05$.

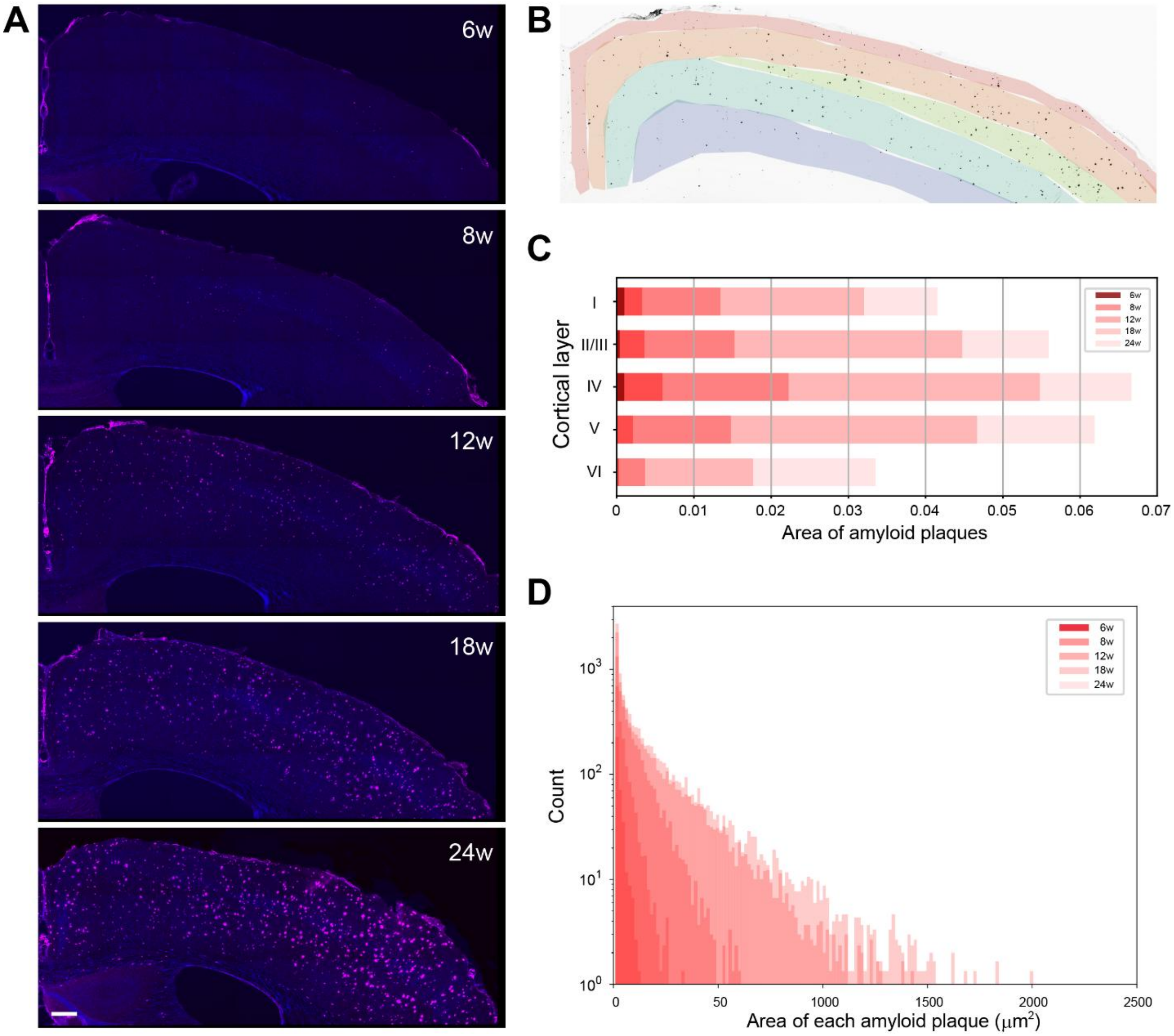


**Fig. S1. Age-dependent progression of amyloid plaque accumulation across cortical layers of male Q-AD mice.** (Related to Fig. 1) (A) Representative coronal brain sections showing age-dependent Aβ deposition in 6–24-week-old male Q-AD mice, visualized by immunostaining with 82E1 antibody. Scale bar: 200 µm. (B) Schematic illustration of cortical layer segmentation used for layer-resolved analysis (L1, L2/3, L4, L5, and L6). (C) Quantification of age-dependent amyloid plaque accumulation in the cerebral cortex of male Q-AD mice. The mean plaque area per animal is shown for each age. (D) Histogram showing the distribution of amyloid plaque sizes at each age. The x-axis represents plaque area (µm²), and the y-axis represents plaque count.

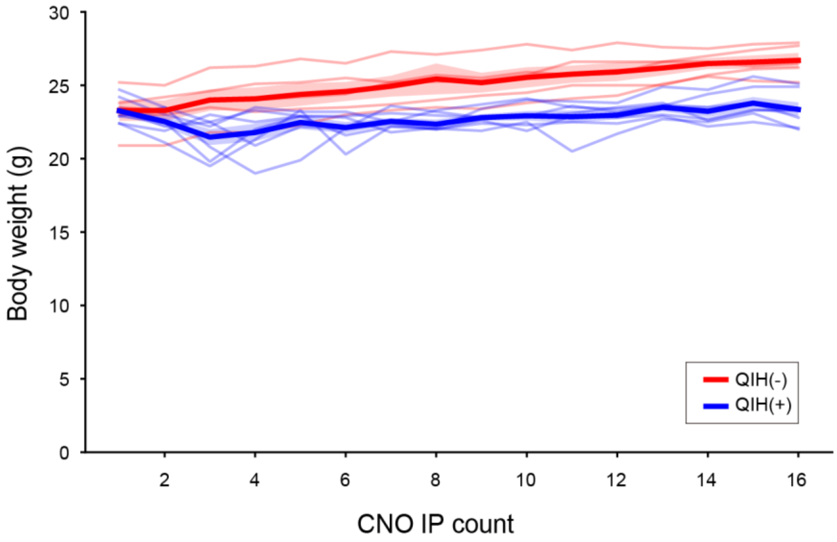


**Fig. S2. Changes in mouse body weight.** (Related to Fig. 1)
QIH(+) mice exhibited a transient decrease in body weight during the early phase of rQIH (first 2–3 injections), followed by a gradual increase over time. Importantly, body weight in QIH(+) mice continued to increase thereafter, although the rate of increase was slightly lower than that in control mice. This early-phase reduction may reflect an acute response to QIH. Thin lines represent individual animals. Thick lines indicate the mean, and shaded areas represent the SEM (QIH(−): n = 5, red; QIH(+): n = 7, blue).

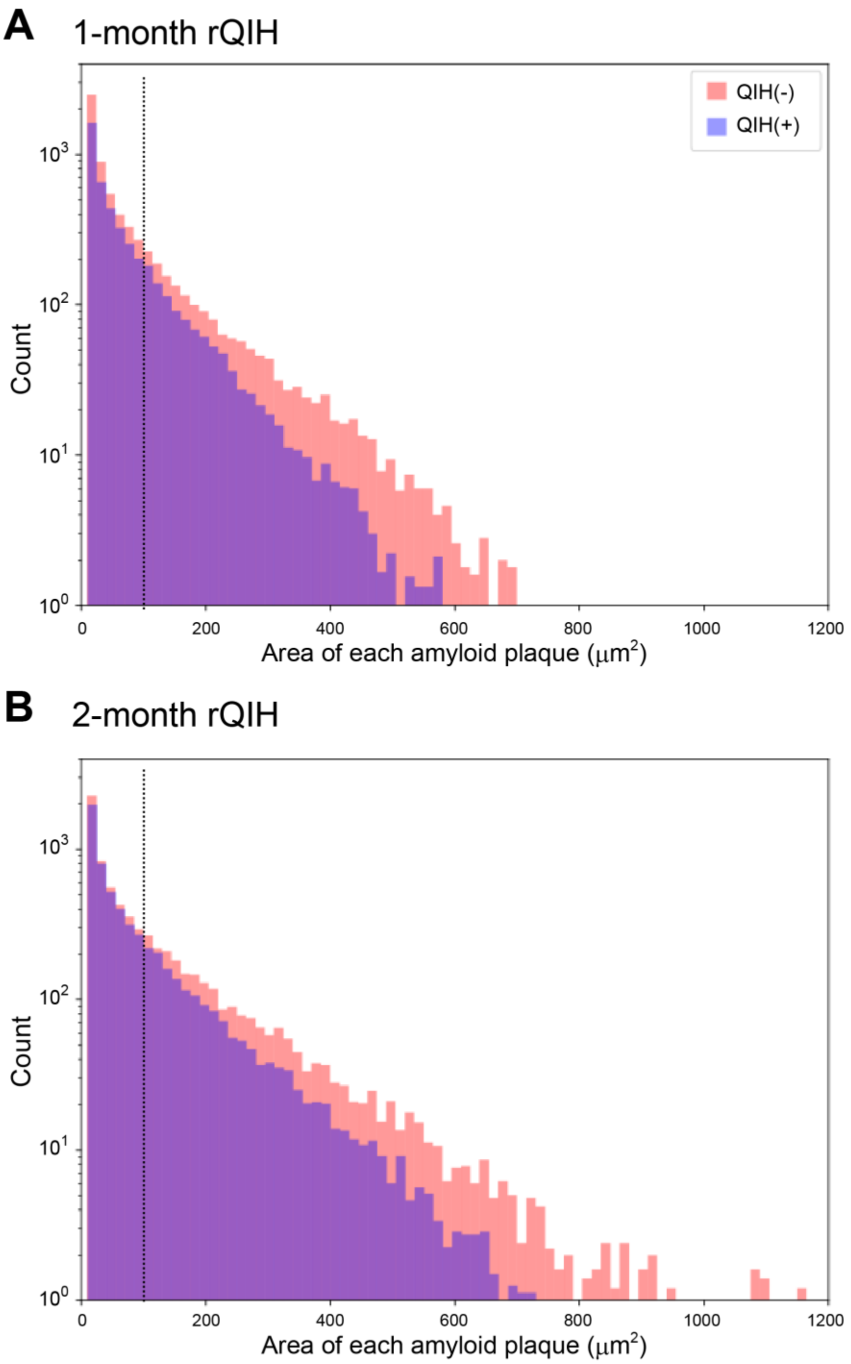


**Fig. S3. Quantification of the effect of rQIH on amyloid pathology.** (Related to Fig. 2) Histogram showing plaque counts as a function of plaque area (A: 1-month rQIH, B: 2-month rQIH). The black dashed line indicates an area of 100 μm². Plaque counts for small plaques were comparable between QIH(+) and QIH(−) groups, whereas the number of large plaques (>100 μm²) was markedly reduced in the QIH(+) group.

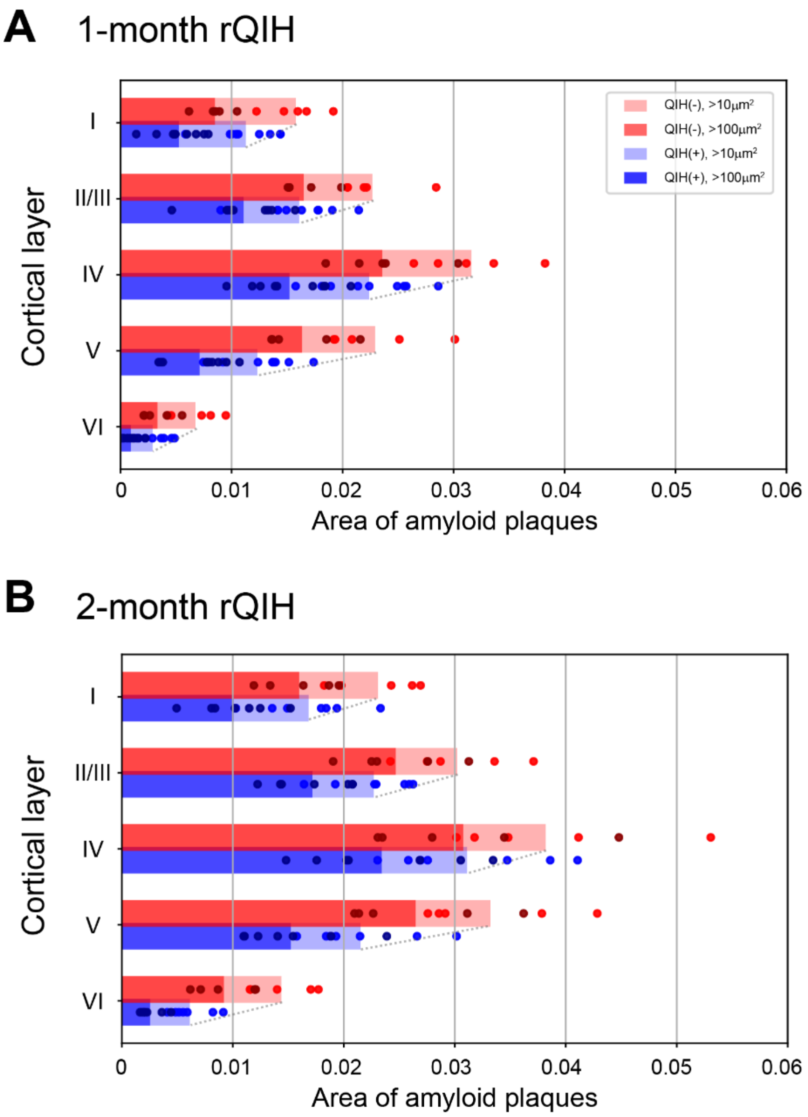


**Fig. S4. Layer-resolved quantification of Aβ deposition in male Q-AD mice subjected to 1-month (A) rQIH or 2-month (B) rQIH.** (Related to Fig. 2)

Data are shown as mean values across individual mice. Statistical analysis was performed using a linear mixed-effects model with Condition (QIH(−) vs QIH(+)) and Layer (L1–L6) as fixed effects and mouse identity as a random effect. In the 1-month rQIH group, a significant main effect of Condition was detected ($p = 0.010$), indicating an overall reduction in Aβ levels following rQIH. In addition, a modest but significant Condition × Layer interaction was observed ($p < 0.05$), suggesting layer-dependent differences in the magnitude of Aβ reduction. Notably, post hoc inspection revealed that the suppressive effect of rQIH was most pronounced in layers 4 and 5. In contrast, in the 2-month rQIH group, although a significant main effect of Condition was observed ($p = 0.027$), no significant Condition × Layer interaction was detected ($p > 0.05$), indicating a layer-nonspecific reduction of Aβ across the cortex. Thin gray lines connecting QIH(−) and QIH(+) bars indicate the direction and relative magnitude of group differences across layers.

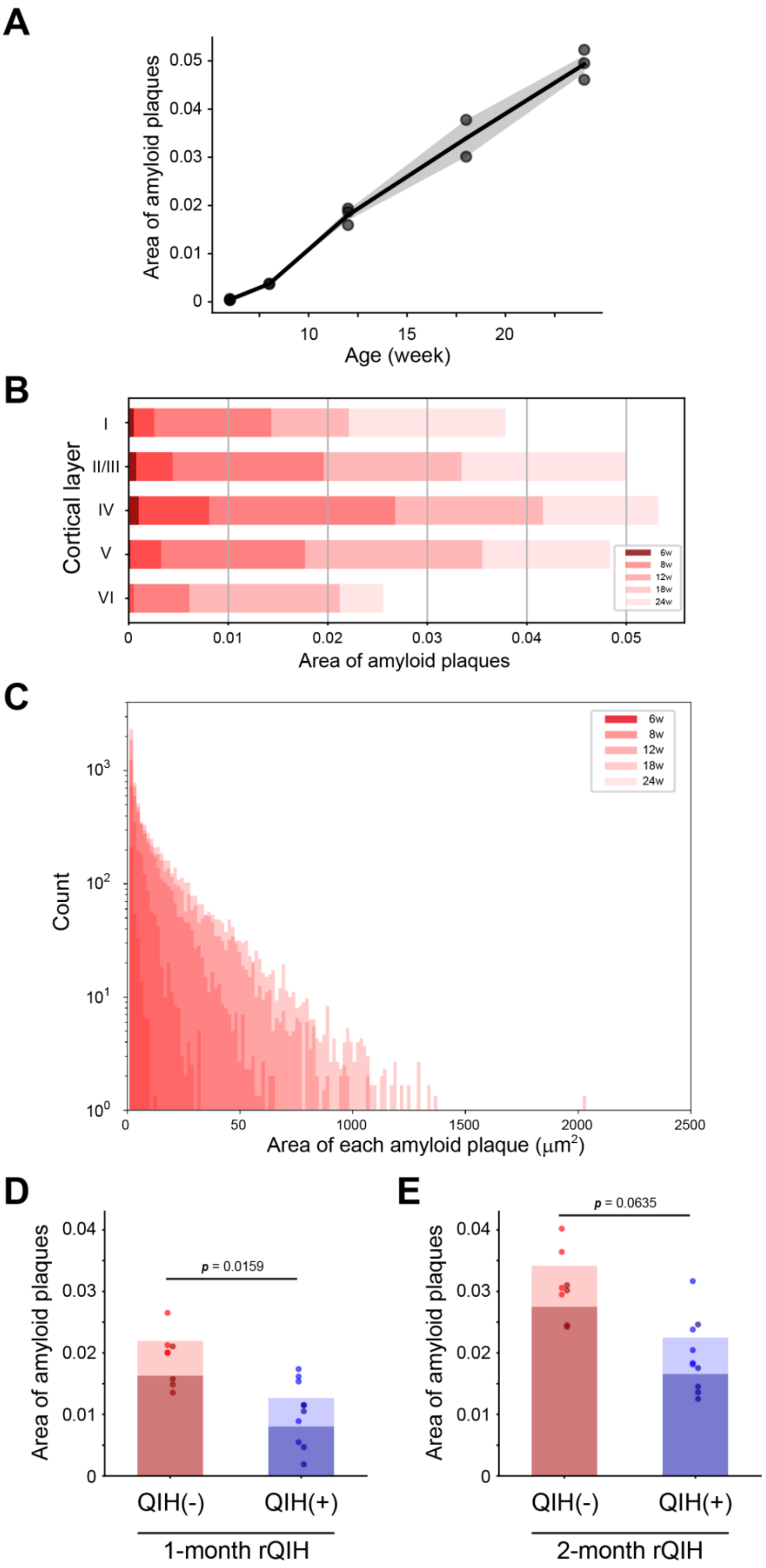
A
Area of amyloid plaques
Age (week)
B
Cortical layer
I
II/III
IV
V
VI
Area of amyloid plaques
6w
8w
12w
18w
24w
C
Count
Area of each amyloid plaque (μm²)
6w
8w
12w
18w
24w
D
Area of amyloid plaques
p = 0.0159
QIH(-)
QIH(+)
1-month rQIH
E
Area of amyloid plaques
p = 0.0635
QIH(-)
QIH(+)
2-month rQIH

**Fig. S5. Quantification of Aβ deposition in female Q-AD mice.** (Related to Figs. 1 and 2) (A) Quantification of Aβ deposition in the motor and somatosensory cortex stained with 82E1 antibody (n = 3 mice per age group, except for 18 weeks, where n = 2). (B) Quantification of age-dependent amyloid plaque accumulation in the cerebral cortex of Q-AD mice. The mean plaque area per animal is shown for each age. (C) Histogram showing the distribution of amyloid plaque sizes at each age. The x-axis represents plaque area ($\mu m^2$), and the y-axis represents plaque count. (D, E) Quantification of Aβ deposition in Q-AD mice subjected to 1-month rQIH (D; QIH(+), n = 5; QIH(-), n = 4) or 2-month rQIH (E; QIH(+), n = 4; QIH(-), n = 4). Dark-colored bars indicate quantification of large plaques (>100 $\mu m^2$). Statistical significance was assessed using the Mann–Whitney U test.

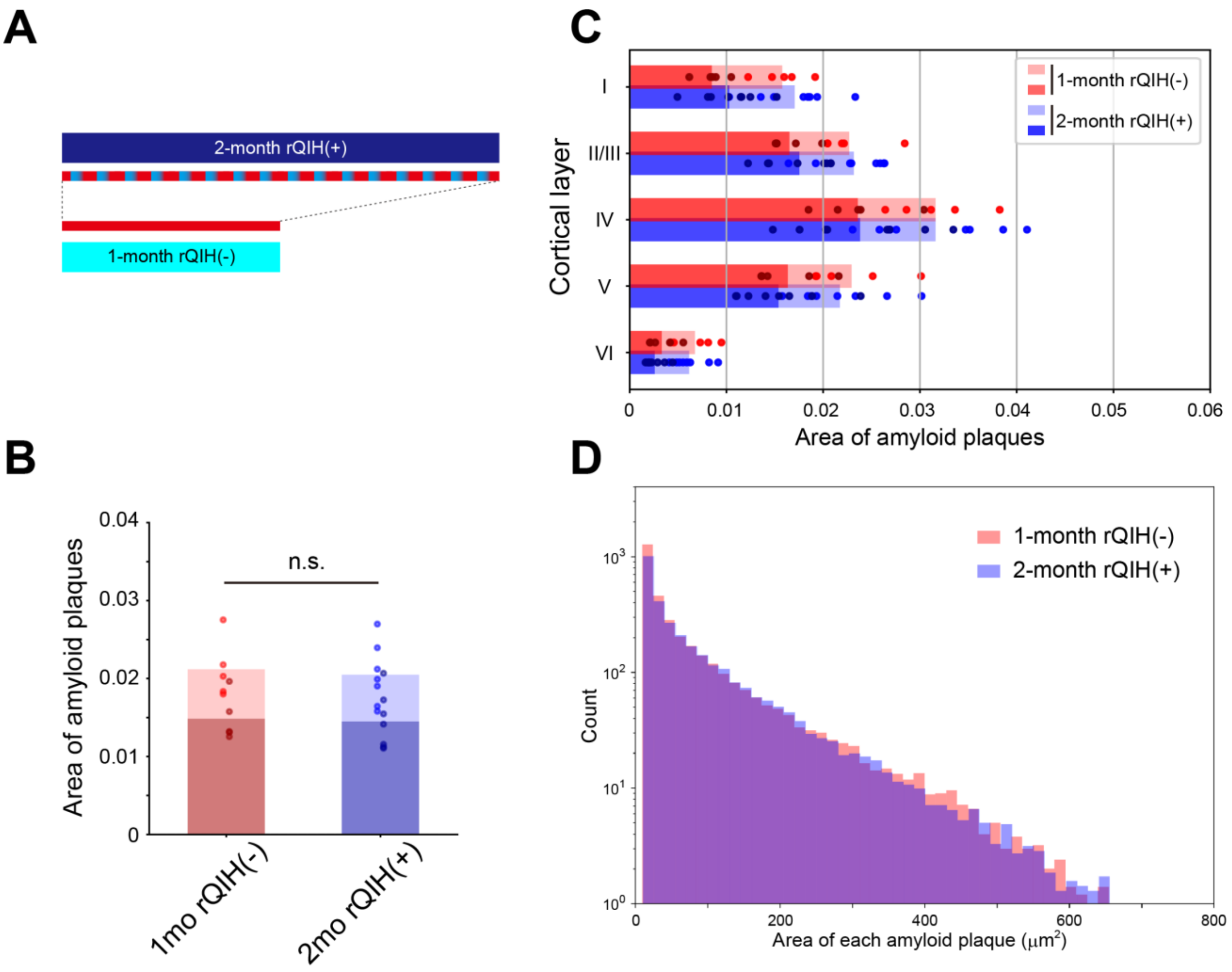


**Fig. S6. Comparison of 1-month rQIH(−) and 2-month rQIH(+).** (Related to Fig. 2) (A) Schematic illustration of the experimental comparison. (B) Quantification of Aβ deposition in male Q-AD mice subjected to 1-month rQIH(−) (n = 5) or 2-month rQIH(+) (n = 7). Dark-colored bars indicate large plaques (>100 μm²). n.s., not significant (Mann–Whitney U test). (C) Layer-resolved quantification of Aβ deposition. Data are shown as mean values per animal. No significant differences were observed between groups. (D) Histogram showing plaque counts as a function of plaque area. The black dashed line indicates an area of 100 μm². Plaque counts across all size fractions were comparable between the 1-month rQIH(−) and 2-month rQIH(+) groups.

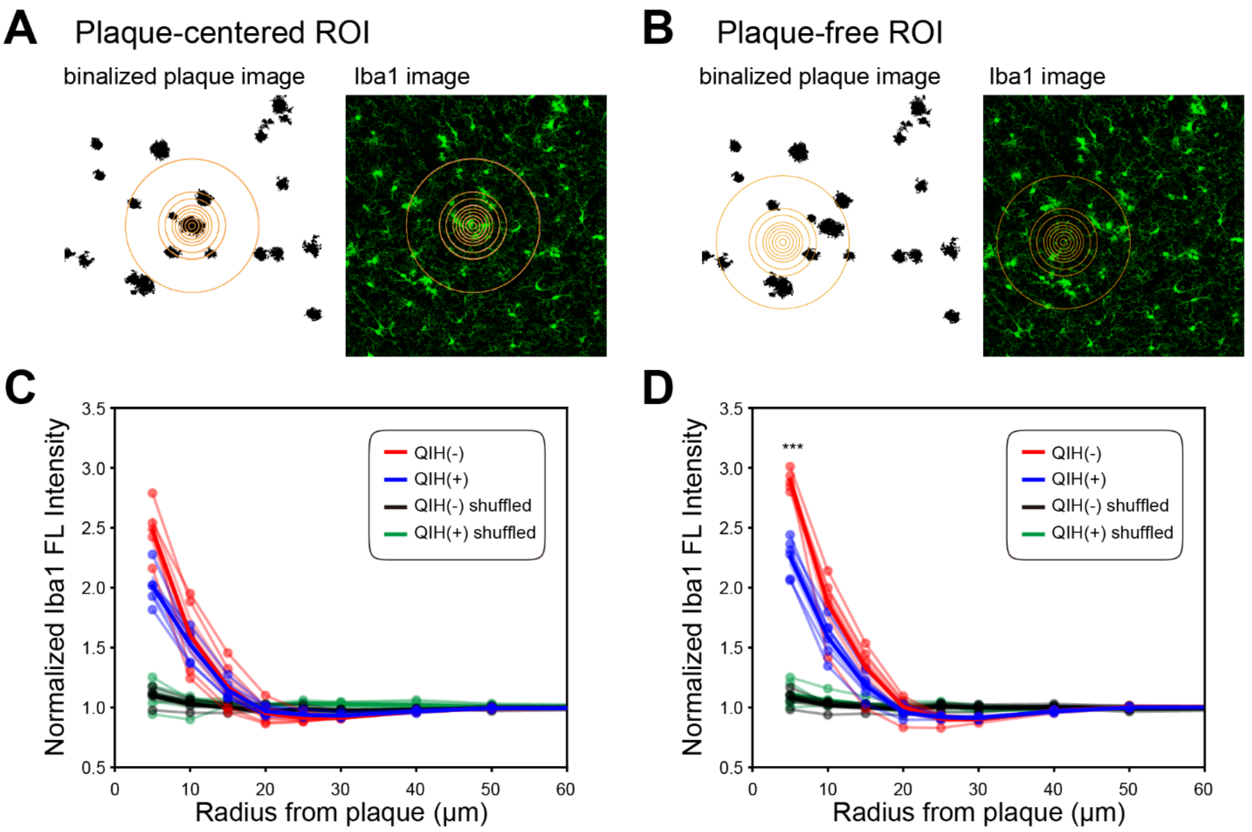


**Fig. S7. Spatial control analysis for plaque-centered ROI quantification.** (Related to Fig. 3) (A) Schematic illustration of plaque-centered concentric circular regions of interest (ROIs). (B) Schematic illustration of plaque-free ROIs used for spatial control (shuffled ROIs). (C, D) Quantification of Iba1 fluorescence intensity as a function of distance from the ROI center for 1-month (C) and 2-month (D) conditions. Concentric circular ROIs were defined either around individual plaques (plaque-centered) or in plaque-free regions (shuffled control), and Iba1 signal intensity was measured radially. The x-axis represents the distance from the ROI center (μm). Red, QIH(−); blue, QIH(+); black, QIH(−) shuffled; green, QIH(+) shuffled. Thin lines represent individual animals, thick lines indicate the mean, and shaded areas represent the SEM. Shuffled ROIs exhibited relatively flat intensity profiles across distances in both groups.
